# Reducing tinnitus via inhibitory influence of the sensorimotor system on auditory cortical activity

**DOI:** 10.1101/2023.06.28.546718

**Authors:** Anne Schmitt, Cora Kim, Stefan Rampp, Elisabeth Bergherr, Michael Buchfelder, Oliver Schnell, Nadia Müller-Voggel

## Abstract

Tinnitus is the subjective perception of a sound in absence of corresponding external acoustic stimuli. Research highlights the influence of the sensorimotor system on tinnitus perception. Associated neuronal processes, however, are insufficiently understood and it remains unclear how and at which hierarchical level the sensorimotor system interacts with the tinnitus-processing auditory system. We therefore asked 23 patients suffering from chronic tinnitus (11 males) to perform specific exercises, aimed at relaxing or tensing the jaw area, which temporarily modulated tinnitus perception. Associated neuronal processes were assessed using Magnetencephalography. Results show that chronic tinnitus patients experienced their tinnitus as weaker and less annoying after completion of relaxing compared to tensing exercises. Furthermore, (1) sensorimotor alpha power and alpha-band connectivity directed from the somatosensory to the auditory cortex increased, and (2) gamma power in the auditory cortex, reduced, which (3) related to reduced tinnitus annoyance perception on a trial-by-trial basis in the relaxed state. No effects were revealed for 23 control participants without tinnitus (6 males) performing the same experiment. We conclude that the increase in directed alpha-band connectivity from somatosensory to auditory cortex is most likely reflecting the transmission of inhibition from somatosensory to auditory cortex during relaxation, where concurrently tinnitus-related gamma power reduces. We suggest that revealed neuronal processes are transferable to other tinnitus modulating systems beyond the sensorimotor one that are e.g. involved in attentional or emotional tinnitus modulation and provide deeper mechanistic insights into how and through which channels phantom sound perception might be modulated on a neuronal level.

**Significance Statement:** Tinnitus describes the perception of auditory phantom sounds. Research suggests that the sensorimotor system impacts on tinnitus perception, associated neuronal mechanisms, however, have remained unclear. Here, chronic tinnitus patients performed exercises with the jaw temporarily reducing (versus increasing) tinnitus perception. Tinnitus reduction was accompanied by an increase of alpha-band connectivity directed from the somatosensory to the auditory cortex and gamma power reduction in the auditory cortex. We suggest that the increase in alpha-band connectivity, when tinnitus is reduced, reflects the transmission of inhibition from somatosensory to auditory cortex, where, in parallel, probably tinnitus related, gamma power reduces. The findings have important implications both for the understanding of phantom sound perception and, more generally, of top-down modulation in healthy and impaired cognition.

## Introduction

Tinnitus is the subjective perception of a sound without a physical sound source (Lockwood, Salvi, and Burkard 2002) affecting about 14% of the population in western societies (Jarach et al. 2022). So far, there is no effective treatment eliminating tinnitus, mostly because processes generating and maintaining tinnitus in the brain are insufficiently understood (Demopoulos et al. 2020). Research showed that tinnitus is initially caused by hearing loss, typically elicited by ageing processes or noise exposure (Kim et al. 2015; Nondahl et al. 2011; Eggermont and Roberts 2004; Reisinger et al. 2023) triggering various neuroplastic changes along the ascending auditory pathway (Haider et al. 2018; Baguley, McFerran, and Hall 2013; Shore, Roberts, and Langguth 2016). Among them, literature emphasizes alterations in oscillatory activity in the auditory cortex of tinnitus patients compared to controls, such as a reduction of alpha and an increase of delta, theta and gamma activity (Adjamian et al. 2012; Kahlbrock and Weisz 2008; Lau et al. 2018; Llinas et al. 1999; Schlee et al. 2014; Weisz et al. 2007; Weisz, Moratti, et al. 2005). The relevance of auditory gamma power is further emphasized by studies relating increased tinnitus loudness to elevated gamma power in the auditory cortex (van der Loo et al. 2009; Sedley et al. 2012; Balkenhol, Wallhausser-Franke, and Delb 2013).

Beyond the abnormalities within the auditory pathway, multiple non-auditory networks are involved in tinnitus most likely reflecting cognitive and emotional processes (Shore, Roberts, and Langguth 2016; Vanneste and De Ridder 2012; Rauschecker, Leaver, and Muhlau 2010; Sedley et al. 2015). Vanneste and de Ridder (2012) postulate a system composed of various subnetworks consisting of different brain areas that are each responsible for different aspects of tinnitus perception.

Recent research has highlighted sensorimotor processes associated with tinnitus (Michiels 2023). A large number of patients can modulate their tinnitus by various movements of the head and neck (Pinchoff et al. 1998; Levine 1999; Levine, Abel, and Cheng 2003), especially in relation with dysfunctions in these regions (Ostermann et al. 2016; Ralli et al. 2016; Algieri et al. 2017). Research indicates that there is a connection between functional disorders of the cervical spine, the jaw joints or musculoskeletal structures and ringing in the ears (Algieri et al. 2017). Beyond, also in ‘normal’ tinnitus, modulation of tinnitus by pressure points or head and neck movements is possible (Levine, Abel, and Cheng 2003; Pinchoff et al. 1998) and patients profit from muscle contraction-based exercises (Sanchez et al. 2007; Michiels 2023). On a neuronal level, sensorimotor processes related to tinnitus have mainly been allocated to subcortical levels of the auditory pathway, such as the cochlear nucleus, inferior colliculus and the trigeminal ganglion and linked to bottom-up processing (Lanting et al. 2010; Skog et al. 2019; Langguth, Elgoyhen, and Cederroth 2019; Marks et al. 2018; Shore 2011; Schilling et al. 2021). Beyond, research indicates that direct connections between the primary somatosensory and primary auditory cortex exist (Lohse et al. 2022), that could be crucial for sensorimotor influence on tinnitus perception. A potential role of the sensorimotor cortex in tinnitus is supported by research showing that the somatosensory cortex integrates somatosensory and auditory input in tinnitus patients (Lanting et al. 2010), exhibits increased functional connectivity compared to controls (Maudoux et al. 2012), and reduces its functional connectivity with the auditory cortex after transcranial direct current stimulation of the auditory cortex (Minami et al. 2015). Moreover, it has been revealed that oral-facial movements increase cerebral blood flow in the sensorimotor cortex of tinnitus patients and concurrently tinnitus loudness (Lockwood et al. 1998) suggesting a potential direct influence of the sensorimotor cortex on tinnitus processing, most likely happening in the auditory cortex. If, however, on a neuronal level, such a direct functional connection between the sensorimotor and auditory cortex responsible for top-down modulation of auditory phantom perception exists, has still to be solved and is aim of the present study.

Given the previously found impact of jaw or orofacial muscle contraction on tinnitus perception (Algieri et al. 2017; Levine, Abel, and Cheng 2003; Michiels 2023; Sanchez et al. 2007), we hypothesize that specific exercises, aimed at relaxing or tensing the jaw area, affect tinnitus perception, and that neurophysiological correlates of this effect become evident in modulations of oscillatory activity in the sensorimotor and auditory cortex and their interaction. Precisely, we hypothesize that relaxing compared to tensing jaw exercises are associated with (1) a reduction of tinnitus loudness and annoyance after completion of exercises, (2) an inhibitory influence directed from sensorimotor to auditory cortex and (3) a reduction of tinnitus-related oscillatory activity in the auditory cortex.

## Materials and Methods

### Participants

Twenty-seven right-handed (one bilateral) volunteers with tinnitus (mean duration 7.9 ± 6.8 years, range six months to 22 years) and twenty-three volunteers without tinnitus participated in the current study. Control participants were age-matched to each tinnitus participant by the +/- 3 years criterion for all but two participants (difference +5 years each). No differences were shown for age between samples (t=0.46, p=0.65). The measurements took place between October 2019 and July 2023 in the laboratory for magnetoencephalography at the Universitätsklinikum Erlangen (Germany). Participants were recruited via flyers posted online on social media. Inclusion criteria for the tinnitus patients were permanent chronic tinnitus (> six months), while the participants for the control group confirmed to not perceive any kind of tinnitus. Further exclusion criteria for both groups were previous neurological diseases and metal in the body that could not be temporarily removed. Four tinnitus patients had to be excluded due to an excessive amount of artefacts in the MEG data (*n* = 4). The remaining *N* = 23 participants with tinnitus (eleven males) had a mean age of 38.5 ± 12.2 years (range 24-61 years). 17 participants perceived their tinnitus bilaterally - four reported left-sided tinnitus and two right-sided tinnitus. Tinnitus characteristics were assessed with an interview (TSCHQ, (Langguth et al. 2007). 19 of 23 participants described their tinnitus as sound and four as noise. Eight patients stated that movements of the jaw, head or neck could influence their tinnitus, the remaining 15 participants negated this question. The N=23 participants without tinnitus (six males) had a mean age of 36.6 ± 11.8 years (range 22-62 years).

All patients were informed about the content of the study, gave their written informed consent prior to taking part in the study and were paid 10 Euro per hour after participation. The Ethics Committee of the Department of Medicine, University of Erlangen-Nürnberg, approved the study (registration number 52_17B).

### Experimental Procedure

The experiment described here was one part of an experimental setting with three different tasks in three consecutive blocks (modulation of tinnitus by jaw exercises, continuity illusion during external tinnitus-like sounds with gaps, modulation of external tinnitus-like sounds by jaw exercises). The present study focuses on the first part of the experiment, where participants were asked to perform relaxing versus tensing exercises with their jaw during tinnitus perception, while perceptual data from the third experiment (same as first part but external tones instead of tinnitus) will also be reported for comparison. In the following, we will describe only the parts relevant for the current study.

Upon arrival in the MEG lab, participants were informed about the study and gave their written informed consent. After that head position coils (HPI) were placed on participants’ heads. The individual head shapes of the participants were collected with a 3D-digital pen. This pen was also used to mark five anatomical points (nasion, LPA, RPA, Cz, inion) as references for later co-registration for source reconstruction. After that, participants were placed in supine position for the main MEG experiment. In the experimental task described here, participants were instructed to complete relaxing (e.g. ‘relax your jaw and let it go as loose as possible’, ‘move your jaw back and forth very gently’) or tensing (e.g. ‘grit your teeth as hard as you can’, ‘imagine you have a very thin object between your teeth and you have to hold it tightly. Under no circumstances should it fall out’) exercises with their jaw for 90 seconds. These exercises elicited the sensation of a relaxed or tensed-up jaw emitting also to the head and neck. Participants were observed with a video camera throughout the entire experiment so that the experimenter could immediately verify whether the participants performed the exercises as they were instructed to. After completing the exercises of the first condition (relaxed or tensed, counterbalanced between participants), participants fixated a black cross in the middle of the screen for three minutes while their brain activity was measured with MEG. In addition, participants were asked to rate perceived loudness and annoyance of their tinnitus on visual analogue scales every minute (loudness scale ranged from 0 to 1 and annoyance scale ranged from −1 to 1). After a short break, the second block started (again exercises, fixation cross and rating). Procedure was the same for the third part with the difference that participants heard after each condition four external tones for one minute respectively (randomised order). These tones were: 1) a 4000 Hz sinus tone, 2) a 7000 Hz sinus tone, 3) white noise with an FFT gaussian filter centred at 4000 Hz, which produced a sound similar to crickets at night, and 4) white noise with Chebyshev filter centred at 4000 Hz, which produced a waterfall-like sound. These sounds were designed according to ‘typical sounds perceived by tinnitus patients’ and represented auditory stimuli for a series of experiments within the framework of a larger project on top-down influences on sound perception in normal-hearing participants and patients suffering from tinnitus (German Research Foundation DFG, project number: 334628700). After each tone participants rated loudness and annoyance of the tone. These tones were adjusted to 55 dB above threshold for each ear and were thus perceived as equally loud in both ears during the experiment. The control group underwent the same procedure with the difference that they did not hear a tinnitus sound during the MEG measurement and consequently were not asked to rate tinnitus loudness and annoyance.

### Behavioural Data Analyses

Behavioural data was analysed with RStudio version 2023.09.1+ (R Core Team 2023, https://www.R-project.org/). Analysis was based on three loudness and three annoyance values per subject and condition with a total of 11 missing loudness values (8%) and 15 missing annoyance values (11%) due to technical problems with the response pads. To examine whether there are differences in subjective tinnitus perception (and perception of external tones as a control condition) between conditions, linear mixed effects models were used (Pinheiro and Bates 2000). Therefore, the main effects of condition (tension/relaxation) and time and their interaction were analysed separately on loudness and annoyance. Participants were integrated as a random effect factor. We tested two models, one with and one without a time factor and took the model with lowest Akaike information criterion (AIC). Analysis showed that time has no significant influence on loudness (*AIC* with time factor: 169.87, *P* > .05; *AIC* without time factor: 155.31) and annoyance (*AIC* with time factor: 122.12, *P* > .05; *AIC* without time factor: 101.17), so we used the model without time factor. Since we had high interpersonal variation, we normalised the data for presentation. Therefore, we divided loudness values by the individual mean loudness value across conditions. For normalisation of annoyance values, we first added a value of one (to get positive values only), divided then by the mean and subtracted then a value of one. After that, we calculated means for each participant and condition over time (as time had no influence). Boxplots were created to display the results. The results were interpreted as significant from a p-value of *P* < .05.

### Data Acquisition

The MEG recordings were accomplished with a 248-channel whole-head-system (Magnes 3600 WH; 4D-Neuroimaging, San Diego, CA, USA) in a magnetically shielded chamber (Vacuumschmelze GmbH, Hanau, HE, Germany). Data were high-pass filtered online at 1 Hz and recorded with a sampling rate of 678.17 Hz. The presentation of visual stimulus material during the MEG recording was controlled using Psychopy version 3 (Peirce 2007) and delivered through a projector via a mirror system.

Structural MRIs were acquired using a high-resolution 3 T MRI-System (Siemens Magnetom Trio, Department of Neuroradiology, Universitätsklinikum Erlangen, Germany).

### MEG Data Analysis

MEG data was analysed with the ‘MNE-Python’ software version v0.24.0 (https://mne.tools/stable/index.html, (Gramfort et al. 2013). The data was first preprocessed. After that, we assessed differences in oscillatory activity between conditions on sensor level using a cluster-based permutation test. Significant clusters were then localised to the brain with a beamformer approach and connectivity between the revealed regions (sensorimotor and auditory cortex) estimated using Partial Directed Coherence. Connectivity analysis was performed using Matlab (The MathWorks, Natick, MA, R 2017b) and the Fieldtrip toolbox (20210330, (Oostenveld et al. 2011). The separate analyses steps are explained in more detail below.

#### Preprocessing

The raw continuous data were segmented into 2-s epochs and notch filtered at 50 Hz and harmonics to eliminate line noise. Furthermore, data sets were down-sampled to 200 Hz. Trials including any visible artefacts due to electromyographic (EMG) noise or technical disturbances were removed through visual artefact rejection. To identify and remove blinks and heartbeat related artefacts we performed an independent component analysis (ICA) using the ‘Picard’ algorithm (Ablin, Cardoso, and Gramfort 2018). Components were visually inspected and artefact-components removed. Furthermore, a reference channel ICA noise cleaning method was applied to remove sources of magnetic noise that are intermittent, and therefore not adequately compensated for by standard, online reference channel correction (Hanna, Kim, and Muller-Voggel 2020).

#### Sensor Level Analyses

First, we aimed at identifying changes in neuronal oscillatory power related to tinnitus reduction after relaxing versus tensing jaw exercises. Activity was contrasted for the relaxed versus tensed state on sensor level. Therefore, power spectral density (PSD) from the epochs of both conditions was computed using multitapers (Percival and Walden 1993). Sensor power was calculated from 1 to 90 Hz in steps of 1 Hz and for both conditions separately (relaxation/tension). Then separate cluster based permutation tests were performed for low (1-40 Hz) and high (40-90 Hz) frequencies to identify the frequencies and channels exhibiting a condition contrast (Maris and Oostenveld 2007). Significant clusters (*P* < .05) in the corresponding frequency bands were then projected to source space.

#### Source Spectral Power Analyses

The generators of sensor effects in source space were assessed with a beamformer approach using the frequency-domain adaptive spatial filtering algorithm Dynamic Imaging of Coherent Sources (DICS, (Gross et al. 2001). Volume conduction models were computed using the FreeSurfer Toolbox (https://surfer.nmr.mgh.harvard.edu/) and the individual MRIs of the participants. Then the surfaces of the inner and outer skull and the scalp were derived and boundary element models (BEM) were constructed using FreeSurfer’s watershed algorithm (Segonne et al. 2004). In a next step, we did a ‘co-registration’ to transfer MEG data and sensor locations in a common coordinate system. For participants without an individual MRI (*n* = 10) we used a standard brain template (‘fsaverage’) for co-registration. After that, we created a surface-based source space for each participant and calculated the forward model. Thereafter, we calculated the cross spectral density matrix (CSD), using morlet wavelets (number of cycles were frequency dependent to optimise frequency over time resolution and varied between 5-15), for the concatenated conditions (for filter calculation) and both conditions separately. According to the significant clusters retrieved on sensor level (7-14 Hz and 32-89 Hz), we selected an alpha frequency range of 7-14 Hz and a gamma frequency range of 32-89 Hz. From the forward models and the CSDs for the concatenated conditions we estimated common spatial filters using DICS for each frequency band (Gross et al. 2001). We then applied the filters for each frequency band to the data of each condition separately to obtain source power values for the two frequency bands and conditions. To quantify the mean difference in alpha and gamma activity between the relaxed and tensed condition we calculated mean change for each frequency by subtracting the average power of the strained from that of the relaxed condition and dividing the result by the former.

In order to focus more precisely on the oscillatory processes taking place within the sensorimotor and auditory regions we added a region of interest (ROI) analysis. Cortical parcellations were derived from the Destrieux cortical atlas (aparc.a2009s, (Destrieux et al. 2010). According to the results in source level analysis, we extracted the power values for the alpha and gamma frequency band for the precentral gyrus, the postcentral gyrus, the anterior transverse temporal gyrus (of Heschl), the lateral aspect of the superior temporal gyrus and the middle temporal gyrus (T2), always for the left and right hemisphere separately. Differences between conditions were quantified using paired t-tests for each region and frequency band. P-values were corrected for multiple comparisons using Bonferroni correction.

#### Brain-Behaviour Regression Analyses

In order to investigate if the observed changes in gamma power are functionally relevant for loudness and annoyance perception we tried to predict individual loudness and annoyance ratings by auditory gamma power. We concentrated on the right auditory cortex, as this was the region with the highest contrast according to the ROI analysis for gamma power. In detail, we extracted for each participant mean gamma power from the right anterior transverse temporal gyrus (of Heschl) for six consecutive data periods corresponding to the six individual rating sessions. We decided to include the last three epochs (6 seconds) before rating because we asked during rating for the actual tinnitus perception (‘How loud/annoying is your tinnitus right now?’). Power values were derived according to the analyses described for ROI analysis with the only difference of averaging power for the last three data segments before rating instead of 3 minutes. We thereby received 138 (23*6) mean auditory gamma power values, which were standardized (z-transformation) for further statistics. Finally, we standardized loudness and annoyance values for subsequent statistics. We then defined two linear mixed effects models to analyse the main effects of condition and oscillatory gamma power, and their interaction on loudness and annoyance perception. Differences between participants were accounted for by integrating them as random effect.

#### Connectivity Analyses

In order to investigate if alpha activity in the sensorimotor cortex is indeed modulating neuronal activity in the auditory cortex we calculated alpha-band directed connectivity between the sensorimotor and auditory cortex. Precisely, alpha-band connectivity was calculated between the postcentral gyrus of the left hemisphere and the transverse temporal gyrus of the right hemisphere, as these were the regions with the highest contrast in ROI analysis. Connectivity was calculated using partial directed coherence (PDC, (Baccala and Sameshima 2001). PDC describes directed relationships among time series in the frequency domain and is based on the concept of Granger causality (Baccala and Sameshima 2001; Granger 1969) which infers causality if past values of A can be used to improve predictions of future values of B. PDC is based on multivariate autoregressive (MVAR) modelling. PDC-values are represented in a range from 0 to 1, with 1 indicating that most of the signal in source *B* is caused by the signal from source *A*, while values close to 0 indicate that there is no information flow from source *A* to *B*. We first projected raw time series into source space by multiplying the raw time series for both conditions separately with a common spatial filter. The spatial filter was created using the LCMV beamformer and the concatenated data of both conditions (Van Veen et al. 1997). Thereby we received time series for both conditions and both ROIs (left postcentral gyrus and right transverse temporal gyrus). For these time series a MVAR model was fitted (‘bsmart’, (Cui et al. 2008)). The model order was set to 15, according to previous approaches (Supp et al. 2007; Muller et al. 2015). Then, a Fourier transform was performed on the resulting coefficients of the MVAR model for 7-14 Hz and a time window of two seconds. These Fourier-transformed coefficients were then used to calculate partial directed coherence from the postcentral gyrus of the left hemisphere to the transverse temporal gyrus of the right hemisphere and vice versa (range of PDC values: 0.002 to 0.186). PDC values were tested for statistical differences between relaxing and tensing conditions using one-tailed paired t-tests for 7-14 Hz (in steps of 1 Hz) and for both directions.

## Results

### Modulation Of Tinnitus Perception

Linear mixed effects models show a significant effect of condition on loudness (*AIC* = 155.31, *t* = −4.89, *P* < 0.001) and annoyance perception (*AIC* = 101.17, *t* = −4.71, *P* < 0.001). As hypothesized, participants perceive their tinnitus significantly less loud and less annoying in the relaxed compared to the tensed condition (**Figure 1**). In contrast, we did not find any effect of condition on loudness and annoyance perception of external tones (*P* = 0.62).

**Figure 1.**
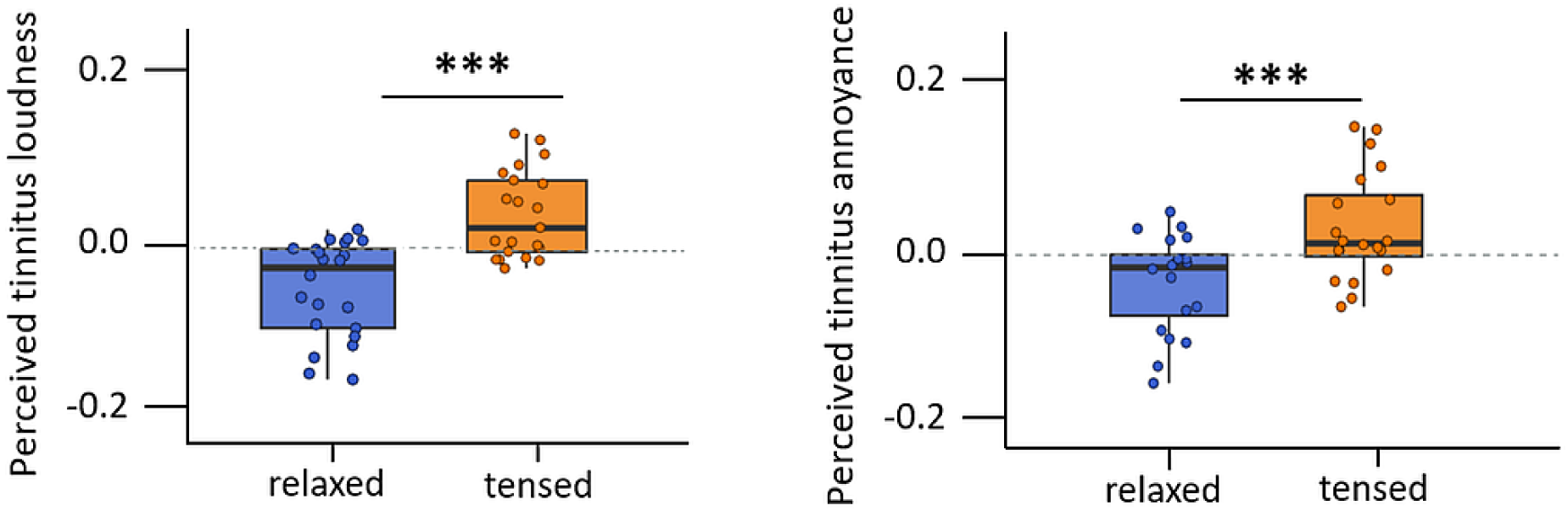
Results of perceptual ratings for the contrast between the relaxed and tensed condition. Left side: Boxplots for the perceived loudness of tinnitus for the relaxed (blue) and tensed (orange) condition. Participants perceived their tinnitus significantly weaker in the relaxed compared to the tensed condition (P < .01**). Right side: Boxplots for the perceived annoyance of tinnitus for a relaxed (blue) compared to tensed (orange) condition. Participants perceived their tinnitus significantly less annoying in the relaxed compared to the tensed condition (P < .001***).

### Modulation Of Oscillatory Power On Sensor Level

Cluster-based permutation tests reveal a significant decrease of power for the relaxed versus tensed condition in the gamma frequency range and peaking at right temporal sensors (**Figure 2A**, 32-89 Hz, cluster *P* < .05) and a significant increase for the relaxed versus tensed condition in the alpha frequency range peaking at fronto-central sensors (**Figure 2B**, 7-14 Hz, cluster *P*< .05). Identical analyses for an age-matched control group did not reveal any significant effect.

**Figure 2.**
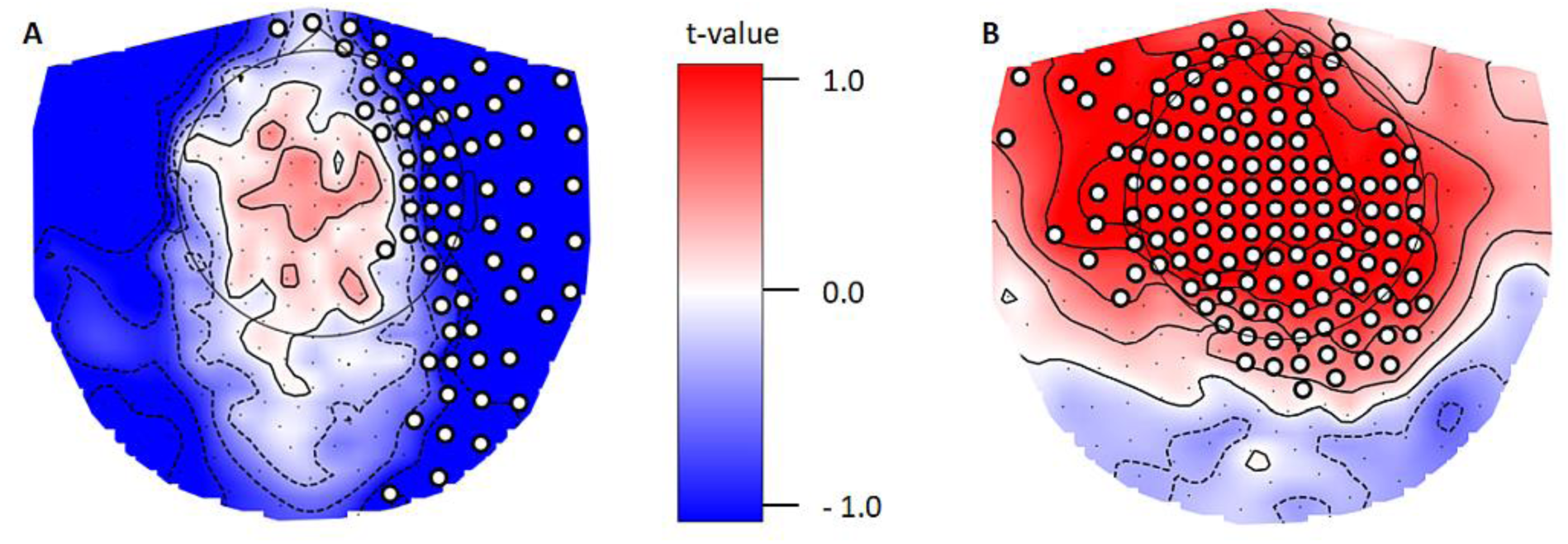
Sensor level cluster-permutation test results for the contrast between relaxed versus tensed states for high (A) and low (B) oscillatory activity. (A) Significant negative cluster in the frequency range of 32-89 Hz (cluster P < .05) peaking at right lateralized sensors. (B) Significant positive cluster for 7-14 Hz (cluster P < .05) for a group of bilateral central sensors. The white dots represent the sensor positions, which form together a significant cluster.

### Modulation Of Oscillatory Power On Source Level

Source results indicate that oscillatory activity in the gamma frequency range (32-89 Hz) is reduced in regions of the superior temporal, transverse temporal and middle temporal gyrus of the right hemisphere (**Figure 3A**) for the relaxed versus tensed condition. Alpha power (7-14 Hz), in contrast, is increased for the relaxed versus tensed condition and this effect is most pronounced in somatosensory and motor cortex in the left and less pronounced in homologous areas of the right hemisphere (**Figure 3B**). We could not reveal any significant power modulations between conditions for the age-matched control group (condition differences for the control group show a somewhat similar pattern, however with only 10% of strength compared to tinnitus group).

**Figure 3.**
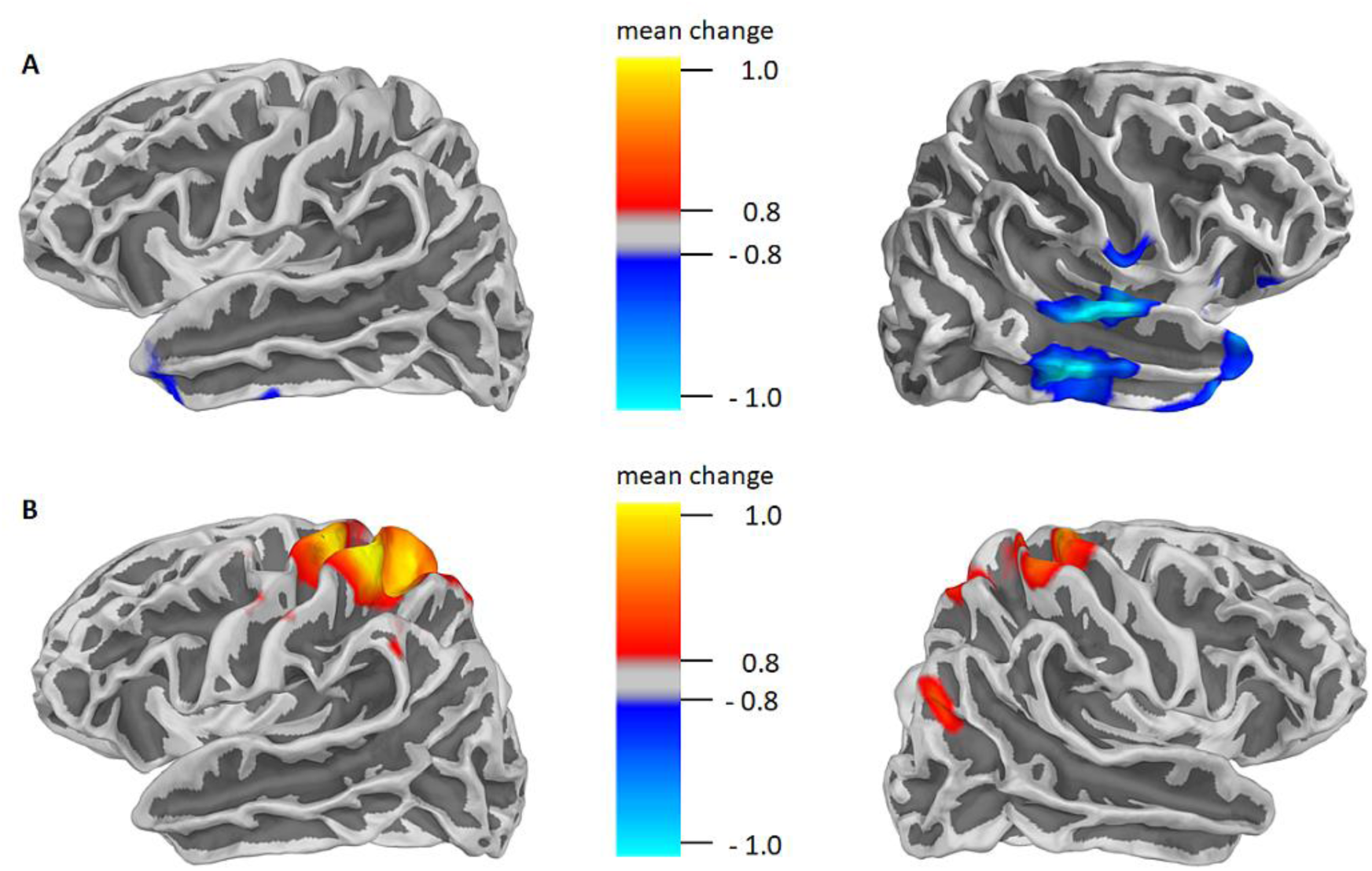
Source power contrast for the relaxed versus tensed condition for gamma (32-89 Hz, A) and alpha (7-14 Hz, B) frequency bands. Mean change is calculated from condition power averages in the form: (P(relaxed) - P(tensed)) / P(tensed). (A) Reduced gamma activity during the relaxed jaw condition in regions of the primary auditory cortex and middle temporal gyrus of the right hemisphere. (B) Increased alpha activity during the relaxed jaw condition in somatosensory and motor cortex of the left hemisphere and a less pronounced increase in homologous areas of the right hemisphere.

The most relevant regions showing significant condition-specific power modulations in the tinnitus group were selected according to a ROI analysis (based on aparc.a2009s atlas, (Destrieux et al. 2010). The extent of power differences between the relaxed versus tensed condition across relevant sensorimotor and auditory regions are described in (**Table 1**). Dependent t-tests show significant differences in the precentral and postcentral gyri of both hemispheres for alpha activity. Alpha activity was significantly higher in the relaxed compared to the tensed condition. For gamma activity, we found significant results in the right hemisphere for the lateral aspect of the superior temporal gyrus and the anterior transverse temporal gyrus. In these regions, gamma activity was significantly lower in the relaxed versus tensed condition. For differences between the relaxed versus tensed condition we found in alpha power the highest t-value in the left postcentral cortex. For differences in gamma power for the relaxed versus tensed condition we found the strongest contrast for the right anterior transverse temporal gyrus (of Heschl). We picked these two regions for connectivity analysis.

**Table 1.** Differences in alpha power and gamma power for the different ROIs (aparc.a2009s atlas, Destrieux et al., 2010). For the contrast between the relaxed versus strained condition, we found in dependent t-tests significant differences in alpha activity for precentral and postcentral gyri of both hemispheres. Alpha activity was significantly higher in the relaxed versus strained condition. Furthermore, we found for the relaxed compared to the strained condition in dependent t-tests significant differences in gamma activity for the right lateral aspect of the superior temporal gyrus and the right transverse temporal gyrus. Gamma activity was significant lower in the relaxed versus strained condition. Statistically significant (after Bonferroni correction) values are highlighted in bold (*P* < .05).

| **Differences in oscillatory power for the relaxed versus strained condition** |  |  |
| --- | --- | --- |
| **Region** | **t-value** | **p-value (corrected)** |
| **Differences in alpha power for the relaxed versus strained condition** |  |  |
| Right precentral cortex | 3.22 | **0.04** |
| Left precentral cortex | 3.61 | **0.02** |
| Right postcentral cortex | 3.19 | **0.04** |
| Left postcentral cortex | 3.70 | **0.01** |
| Right lateral aspect of the superior temporal gyrus | 1.05 | > 0.99 |
| Left lateral aspect of the superior temporal gyrus | 0.85 | > 0.99 |
| Right middle temporal gyrus (T2) | 0.73 | > 0.99 |
| Left middle temporal gyrus (T2) | 0.44 | > 0.99 |
| Right anterior transverse temporal gyrus (of Heschl) | 0.65 | > 0.99 |
| Left anterior transverse temporal gyrus (of Heschl) | 1.05 | > 0.99 |
| **Differences in gamma power for the relaxed versus strained condition** |  |  |
| Right precentral cortex | -0.58 | > 0.99 |
| Left precentral cortex | 1.46 | > 0.99 |
| Right postcentral cortex | -0.90 | > 0.99 |
| Left postcentral cortex | 0.88 | > 0.99 |
| Right lateral aspect of the superior temporal gyrus | -3.16 | **0.05** |
| Left lateral aspect of the superior temporal gyrus | -1.88 | 0.74 |
| Right middle temporal gyrus (T2) | -2.96 | 0.07 |
| Left middle temporal gyrus (T2) | -2.26 | 0.34 |
| Right anterior transverse temporal gyrus (of Heschl) | -3.24 | **0.04** |
| Left anterior transverse temporal gyrus (of Heschl) | -0.11 | > 0.99 |

### Association Between Auditory Gamma Power And Tinnitus

Linear mixed effects models show an interaction effect between condition and auditory gamma power on annoyance perception (*AIC* = 109.21, *t* = 2.25, *P* < 0.05) besides the already described main effect of condition (*P* < 0.001). An increase in gamma power was associated with an increase in tinnitus annoyance, especially in the relaxed condition, while this association was absent in the tensed condition (**Figure 4a**). In contrast, we found no significant effects for the association between auditory gamma power and loudness perception (for description on an explorative level see Figure **4b**).

**Figure 4.**
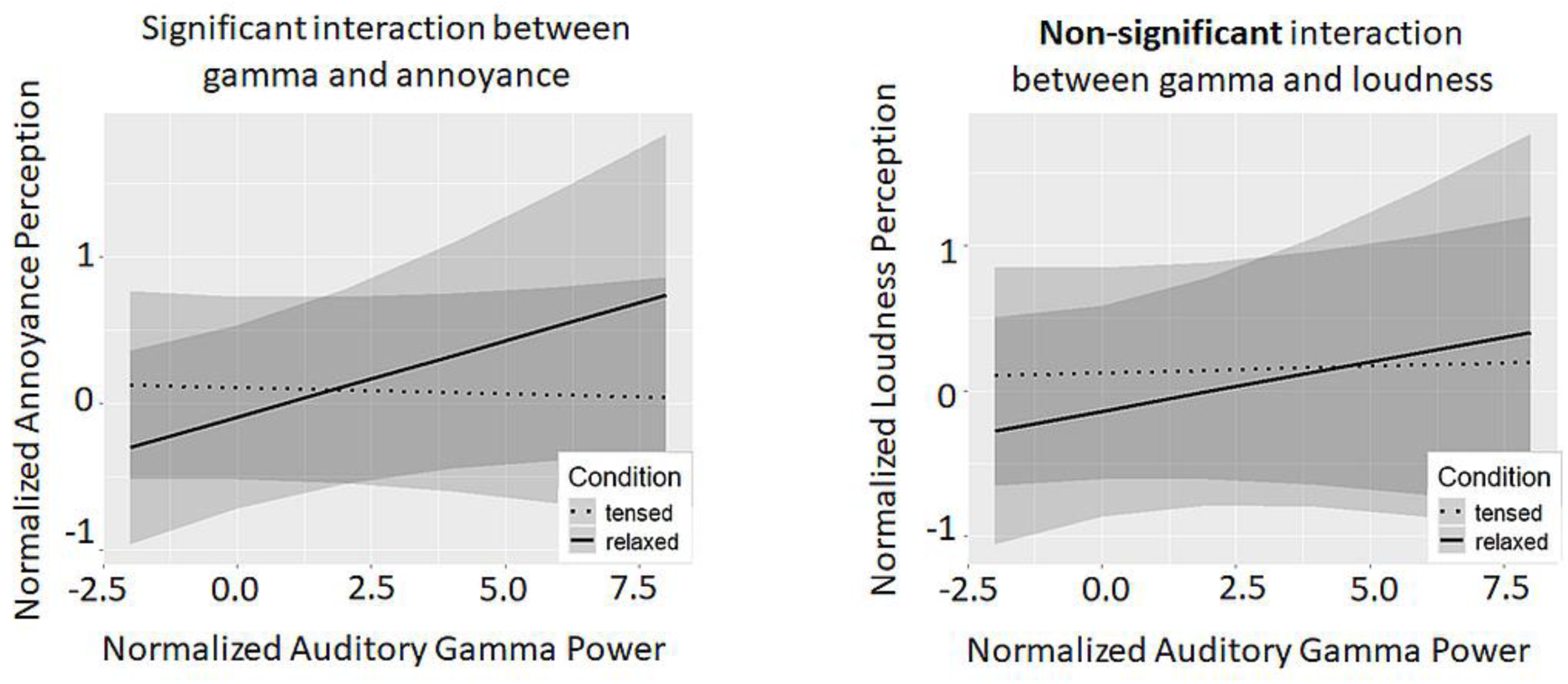
Results for the association between gamma power in the right primary auditory cortex and predicted tinnitus ratings. The left panel shows the significant interaction between auditory gamma power and tinnitus annoyance ratings (P < .05). An increase in auditory gamma power relates to an increase in perceived tinnitus annoyance in the relaxed condition, while this association is absent in the tensed condition. The right panel displays the non-significant interaction between auditory gamma power and tinnitus loudness ratings, which illustrates, on an explorative level, a similar pattern as the interaction between auditory gamma power and tinnitus annoyance. The grey shades indicate the 95% credible intervals.

### Communication Between The Somatosensory And Auditory Cortex

Directed connectivity was analysed from the left postcentral gyrus to the right transverse temporal gyrus for the alpha frequency range (7-14 Hz). As control condition, we calculated connectivity in the opposite direction, from the right transverse temporal gyrus to the left postcentral gyrus. For the relaxed versus tensed condition, connectivity was significantly increased for 11-14 Hz and from the left postcentral gyrus to the right transverse temporal gyrus (*t* = 2.18, *P* < .05, **Figure 5**). Connectivity for the opposite direction, from the right transverse temporal gyrus to the left postcentral gyrus showed no significant differences between conditions. For the age-matched control group we could not reveal any increase in alpha-band connectivity in neither direction.

**Figure 5.**
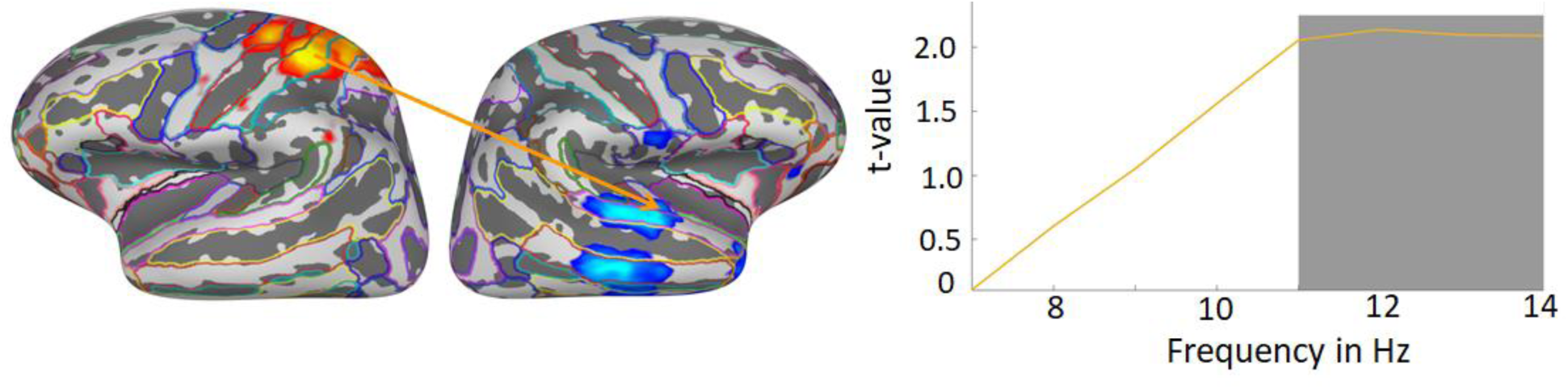
Results for the directed connectivity analysis from the postcentral gyrus of the left hemisphere to the transverse temporal gyrus of the right hemisphere in the alpha frequency range (7-14 Hz). Directed connectivity from the left postcentral to the right transverse temporal cortex increased for 11-14 Hz after relaxing compared to tensing exercises. The significant frequency range is marked in grey (P < .05). Directed connectivity from the right transverse temporal gyrus to the left postcentral gyrus, in contrast, was not modulated for 7-14 Hz.

## Discussion

The present study shows that chronic tinnitus patients perceive their tinnitus significantly reduced after having executed relaxing compared to tensing jaw exercises. On cortical level, sensorimotor alpha power and alpha connectivity from sensory to auditory cortex increases and auditory gamma power reduces after relaxing compared to tensing exercises. Importantly, we could not find any modulations of oscillatory power or connectivity in an age-matched control group, who performed the same experiment, indicating that the observed modulation of neuronal activity relates to interactions with the tinnitus percept.

### Modulation Of tinnitus Perception By The Sensorimotor System

Perceptual data shows that participants perceived their tinnitus significantly weaker and less annoying up to at least three minutes after relaxing compared to tensing exercises. Even though we cannot exclude that the ratings were biased by socially desirable responses having tempted the participants to rate their tinnitus weaker after relaxing exercises, we are confident that the ratings reflect true tinnitus perception due to their association with neuronal modulations. Beyond, the finding of reduced tinnitus perception after relaxing exercises is consistent with research showing that tinnitus patients benefit from relaxing exercises (Sanchez et al. 2007) as part of cognitive behavioural therapy (Schecklmann et al. 2023) or when suffering from temporomandibular disorder (Sanchez and Rocha 2011). Most interestingly, we could not elucidate any differences in loudness and annoyance perception of external sounds after relaxing versus tensing exercises for the same participants. We therefore suggest that the sensorimotor system is not generally inhibiting auditory processing in the sense of a global downregulation of processing but seems to specifically reducing neuronal activity related to tinnitus perception.

## Neuronal Results

### Reduced Tinnitus Perception Is Accompanied By Reduced Auditory Gamma Activity

Relaxing jaw exercises did not only lower tinnitus perception, but reduced gamma activity in auditory brain regions, especially of the right hemisphere, and most pronounced in the transverse temporal gyrus and lateral aspect of the superior temporal gyrus. This is consistent with research showing positive correlations between the level of auditory cortical gamma power and perceived tinnitus intensity (van der Loo et al. 2009; Balkenhol, Wallhausser-Franke, and Delb 2013) or auditory gamma power increases during tinnitus perception compared to when phantom sounds are absent (Ortmann et al. 2011; Weisz et al. 2007). The significant gamma decrease in the lateral aspect of the superior temporal gyrus also matches a reduced processing of tinnitus, since the superior temporal gyrus is associated with auditory processing of sounds (Yi, Leonard, and Chang 2019) and auditory hallucinations (Jaroszynski et al. 2022). Apart from tinnitus, auditory cortical gamma power has been associated with conscious processing of acoustic stimuli or imagery and correlates with spiking activity of the underlying tissue (Cervenka et al. 2013; Hayat et al. 2022; Pantev et al. 1991; Teng et al. 2017; Martin et al. 2018; Lachaux et al. 2012). This corroborates that ongoing decrease of gamma power in auditory cortex relates to reduced tinnitus perception after completion of relaxing compared to tensing jaw exercises. Behavioural relevance of auditory gamma power decrease is further supported by the regression results showing that right auditory gamma power successfully predicts tinnitus annoyance, with the relaxing condition driving the effect. Interestingly, this association is absent for the tensed condition. We suggest that tinnitus patients might have been already in a chronic tensed state (Schecklmann et al. 2023) so that tensing exercises had a significantly lower potential to further intensify ‘tension’ and thereby modulate tinnitus compared to relaxing exercises. Unfortunately, we did not quantify the extent of tension during the experiment or baseline to prove that assumption directly. However, following that reasoning we expect that the variance of tinnitus ratings is lower for the tensed compared to the relaxed condition, which indeed tends to be the case (*F* =1.44, *P* = 0.08). We did not find any significant effect pointing to an association between auditory gamma power and loudness perception, which could be partly due to the little amount of perceptual ratings lowering statistical power (on an explorative level a similar pattern as for annoyance perception is observable). Given the common difficulty of reliably assessing the subjective characteristics of tinnitus (Langguth and De Ridder 2023), there could have been also differences in the capability of judging the different aspects of tinnitus. Further studies, including a significantly higher number of perceptual ratings, will have to clarify how other tinnitus characteristics beyond annoyance are associated with auditory gamma power.

The right-lateralization of the auditory gamma effect is in line with research showing an increase of right auditory gamma activity during acute tinnitus perception (Ortmann et al. 2011), a right hemispheric lateralization associated with tinnitus distress (Weisz, Wienbruch, et al. 2005), and differences in right auditory cortical volume in tinnitus patients compared to healthy controls (Ma et al. 2022). Findings regarding lateralized effects in auditory cortex also relate to the laterality of tinnitus perception (Lockwood et al. 1998; Weisz et al. 2007). Since most of our participants perceived their tinnitus bilaterally, it remains unclear to what extent tinnitus laterality had an influence on the results.

### Increase Of Alpha Activity In The Sensorimotor System After Relaxing Jaw Exercises

After completion of relaxing versus tensing jaw exercises, we found a significant increase of alpha power in the sensorimotor cortex of both hemispheres. The most pronounced alpha power increase during relaxation locates to a region of the somatosensory cortex consistent with the cortical representation of the head/neck area most likely related to corresponding sensation of this area after completion of the exercises. An increase of alpha power within the sensorimotor cortex has been associated with its functional inhibition (Jensen and Mazaheri 2010; Haegens et al. 2011; Stolk et al. 2019). Alhajri and colleagues (2018) revealed that sensorimotor alpha activity decreases during, but also after physical practice. This suggests that the alpha power increase in the current study, measured after completion of the relaxing (compared to tensing) exercises, might reflect a sustained state of relaxation becoming evident in functional inhibition of the head/neck area. This is also in line with a finding from Flüthmann and colleagues (2019) who showed an increase in sensorimotor alpha activity during muscle relaxation, which they relate to corresponding inhibitory processes. In conclusion, we suggest that after relaxing compared to tensing exercises the sensorimotor cortex is in a locally inhibited state. For reducing the actual tinnitus percept (most likely reflected in the reduction of auditory cortical gamma activity), the somatosensory cortex must transfer this inhibition to the auditory cortex.

### Transmitting Inhibition From The Sensorimotor To The Auditory Cortex Through An Increase Of Directed Alpha-Band Connectivity

In parallel to the local sensorimotor alpha power increase after relaxing compared to tensing exercises, we found an increase of alpha-band connectivity directed from somatosensory to auditory cortex most likely indicating the transmission of inhibition to the auditory cortex. Importantly, we found no significant alpha-band connectivity modulation between conditions in the opposite direction, from auditory to somatosensory cortex. That an increase in directed alpha-band connectivity could represent a neuronal mechanism for targeted inhibition between brain regions, especially in a top-down manner with higher order regions asserting influence on lower-level stimulus-processing regions, is supported by increasing evidence, e.g. in attention tasks (Jensen and Mazaheri 2010; Zhao and Wang 2019; Popov, Kastner, and Jensen 2017; Hanna et al. 2024). Additionally, macaque studies highlight the role of alpha/beta connections in top-down modulation of gamma power in sensory cortices (Bastos et al. 2020). This is in line with our results showing an increase of alpha-band connectivity directed from the somatosensory to the auditory cortex, where concurrently auditory gamma power reduces. Importantly, the described neuronal processes take place after completion of the jaw exercises mostly ruling out bottom-up driven interactions between sensorimotor and auditory cortex and emphasizing our interpretation as a top-down process. Our results significantly add to previous tinnitus research having emphasized subcortical processes and bottom-up driven interactions between somatosensory and auditory activity (Lanting et al. 2010; Shore 2011; Schilling et al. 2021; Shore et al. 2008) by depicting how the somatosensory cortex top-down modulates tinnitus-related auditory cortical activity. This knowledge will help in understanding better through which ‘channels’ tinnitus can be modulated and inspire new therapy approaches.

## Conclusion

With the present study, we aimed at describing in detail how the sensorimotor system affects tinnitus processing by tracking cortical neurophysiological changes that accompany a reduction in perceived tinnitus after sensorimotor intervention. We could show that, when tinnitus patients are in a relaxed state elicited by preceding jaw exercises, tinnitus perception decreases. On a neuronal level, the sensorimotor cortex turns into an inhibited state and transfers inhibition to the auditory cortex where gamma power reduces. The trial-by-trial association between auditory gamma power and tinnitus annoyance during relaxation underlines the functional relevance of this process for tinnitus perception. By tracing specific neuronal processes, acting within and between sensorimotor and auditory cortices during tinnitus modulation, we establish an analytical framework that could help tackling and understanding neural mechanisms underlying top-down modulation of tinnitus by other interventions such as e.g. cognitive trainings for relaxation or attention directing (Weise, Kleinstauber, and Andersson 2016) and inform related interventional approaches.

## Data availability

The data that support the findings of this study are available upon request to the corresponding author.

## Acknowledgements

We would like to thank Martin Kaltenhäuser, Maximilian Vollmayer and Antonia Keck for assistance in data collection.

